# Enrichment of Skeletal Stem Cells from Human Bone Marrow Using Spherical Nucleic Acids

**DOI:** 10.1101/2019.12.19.882563

**Authors:** Miguel Xavier, Maria-Eleni Kyriazi, Stuart A. Lanham, Konstantina Alexaki, Afaf H. El-Sagheer, Tom Brown, Antonios G. Kanaras, Richard O.C. Oreffo

## Abstract

Human bone marrow (BM) derived stromal cells contain a population of skeletal stem cells (SSCs), with the capacity to differentiate along the osteogenic, adipogenic and chondrogenic lineages enabling their application to clinical therapies. However, current methods, to isolate and enrich SSCs from human tissues remain, at best, challenging in the absence of a specific SSC marker. Unfortunately, none of the current proposed markers, alone, can isolate a homogenous cell population with the ability to form bone, cartilage, and adipose tissue in humans. Here, we have designed DNA-gold nanoparticles able to identify and sort SSCs displaying specific mRNA signatures. The current approach demonstrates the significant enrichment attained in the isolation of SSCs, with potential therein to enhance our understanding of bone cell biology and translational applications.

**TABLE OF CONTENTS:** 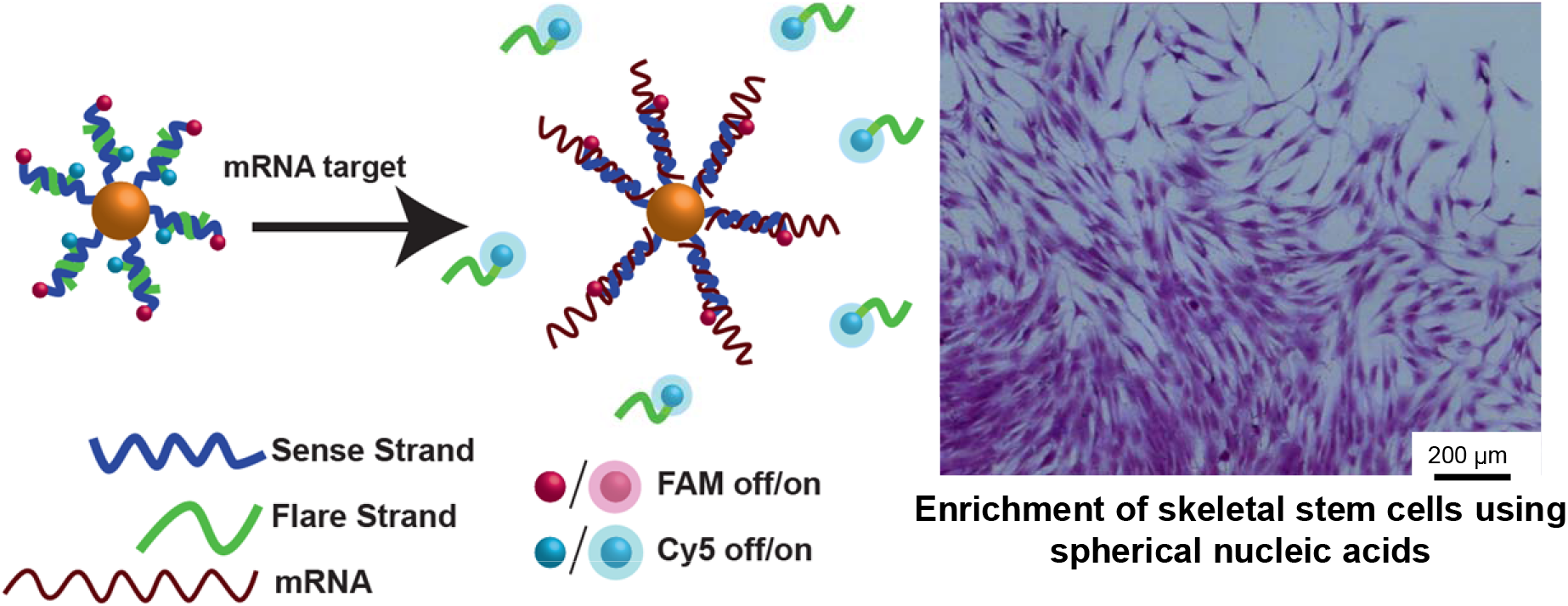

## INTRODUCTION

In the last decade, progress in colloid chemistry has enabled the synthesis of advanced, functional nanoparticles (NPs) that can survive complex biological media and perform programmed tasks including cell targeting, sensing, and drug release.^1, 2^ Mirkin and co-workers were the first to demonstrate how the surface of gold NPs could be functionalised with a shell of synthetic oligonucleotides attached to the gold surface *via* a thiol bond.^3^ This new design of oligonucleotide – coated NPs, also termed spherical nucleic acids (SNAs) have subsequently emerged as live cell probes for the detection of targets including RNA, molecules and ions.^4–12^ The extensive use of SNAs within the biomedical field is primarily due to their high specificity, binding of complementary oligonucleotide strands and improved resistance towards DNA enzymatic degradation as well as the enhanced uptake of SNAs by multiple types of cells.^13, 14^ Seferos *et al.* were the first to show how SNAs could be hybridised to shorter fluorophore bearing oligonucleotides for the targeted detection of mRNA. The authors demonstrated the detection of survivin mRNA, where a significant increase in fluorescence intensity was recorded compared to treatment with non – targeting NPs.^15^ Following this work, several studies have demonstrated the use of SNAs for the detection of mRNA targets with high accuracy and specificity. Lahm *et al.* demonstrated the use of SNAs for the detection of nanog and gdf3 mRNA in embryonic stem cells as well as induced pluripotent stem (iPS) cells of murine, porcine and human origin. Target positive cells were sorted after which nanog – specific probes identified reprogrammed murine iPS cells.^16^ The detection of nanog mRNA was also taken advantage of by Owen *et al.* for the detection and isolation of cancer stem cells.^17^ In contrast, Wang *et al.* developed SNAs to enable self – activating and imaging of mRNA sequential expression (tubb3 and fox 3 mRNA) during the neural stem cell differentiation process.^18^ Recently, we have demonstrated that SNAs can be used to detect vimentin mRNA in skin wounds as vimentin mRNA is expressed during the epithelial to mesenchymal transition that occurs during wound healing.^19^ In addition, we have designed SNAs to detect hymyc1 mRNA, responsible for the regulation of the balance between stem cell self – renewal and differentiation, in real time in live *Hydra Vulgaris*.^20^ We have also shown the development of SNAs and dimer SNAs, which can detect up to two different mRNA targets whilst coordinating the release of two different types of drugs within the same local microenvironment.^21, 22^

Recently, the SNA design has evolved to include mRNA detection directed by an external stimulus. Lin *et al.* demonstrated the use of such a strategy for the specific detection of manganese superoxide dismutase (MnSOD) mRNA. Following application of oligonucleotides modified with a photoactivated linker, mRNA detection was present only in select cells at a desired time point *via* the use of two-photon illumination. By illuminating specific cells of interest, mRNA detection was visualized only in these specific cells, while neighboring cells remained fluorescently dormant.^6^

The design of SNAs has been further exploited for the detection of other targets including miRNAs and, recently, for the delivery of oligonucleotides or peptides to activate the immune system.^5, 23–28^ Zhai *et al.* reported the detection of microRNA – 1246, a breast cancer biomarker, in human plasma exosomes, whereas, Zhao *et al.* used the same design for the direct detection of microRNA – 375.^29^ In contrast, Ferrer *et al.* developed a dual targeting SNA that could coordinate the delivery of a nucleic acid specific for toll like receptor 9 (TLR9) inhibition and a small molecule (TAK – 242) that inhibited TLR4; with the goal of attenuating inflammation by downregulating pro-inflammatory markers downstream of each receptor.^24^ Undoubtedly, SNAs have emerged as a versatile tool, with significant potential within the biomedical field.

An area of research that has come to the fore in recent years is the application of stem cells in regenerative medicine. The ready accessibility of human bone marrow stromal cells (BMSCs) from bone marrow and their ability to differentiate into bone-forming osteoblasts when implanted *in vivo*, has garnered significant scientific interest and driven the application of SSCs in the clinic.^30^ Bone marrow (BM) is the soft connective tissue within bone cavities that functions to produce blood cells and as a store for fat. Traditionally BM has been seen as an organ composed of two main systems: the hematopoietic tissue and the supportive stroma.^31^ Hematopoietic stem cells (HSCs) sustain the generation of all blood cell types and are thus invaluable for the treatment of hematopoietic disorders.^32^ In contrast, the BM stroma contains mesenchymal stem/stromal cells (MSCs). These are defined as unspecialized cells that lack tissue – specific characteristics and maintain an undefined phenotype.^33^ The term MSC was originally coined in reference to a hypothetical common progenitor of a wide range of “mesenchymal” (non-hematopoietic, non-epithelial, mesodermal) tissues. The term represents a heterogeneous cell population when cultured *in vitro* and are likely to include cells with a wide spectrum of regenerative potential from specific terminal cell type progenitors to multipotent stem cells, named SSCs.^34, 35^ The SSC is used here to refer specifically to the self-renewing stem cell of the bone marrow stroma responsible for the regenerative capacity inherent to bone with osteogenic, adipogenic and chondrogenic differentiation potential *in vivo*.^34, 36^ However, a number of challenges limit SSC application including the limited numbers of SSCs in BM aspirates (1 in 10-100,000), and the absence of specific markers limiting facile homogenous isolation of the SSC from human tissue. Thus, there remains limited understanding of SSC fate, immunophenotype and simple selection criteria; all of which have proven to be limiting factors in the widespread clinical application of these cells.^34^

Current techniques employed for SSC sorting are based on fluorescence and magnetic activated cell sorting (respectively named FACS and MACS) combined with plastic culture adherence. However, FACS and MACS depend on the use of antibodies to target antigens present on the cell membrane, in the cytoplasm or the nucleus, with detection of internal antigens requiring the cells to be fixed and therefore unable to be cultured further.^37^ Cells separated *via* FACS depend on the absence or presence of a fluorescence tag, whereas, MACS uses magnetic beads attached to a primary antibody. Tagged cells are thus retained in the device by a strong magnetic field.^34^ The use of both techniques is hindered by the cost and time of sample processing, although, the single largest hurdle remains the lack of specific cell membrane markers for SSCs.^34^ Widely used markers, such as STRO-1, interact with only a small percentage of BM cells that contain a sub-population that includes the SSC/progenitor population. Although, a range of cell surface markers can enrich for SSCs none of the proposed markers, in isolation, can isolate single cells with the ability to form bone, cartilage and adipose tissue in humans and a specific marker for the SSC, to date, remains elusive.^38–40^

In this study, we take advantage of the versatile nature of SNAs not only to detect SSCs based on mRNA expression but also to rapidly isolate SSCs from human BM; a critical requirement for tissue engineering strategies harnessing SSCs to aid bone regeneration. BM cells expressing specific mRNA targets were isolated *via* FACS and subsequently used to isolate SSCs, without affecting long-term skeletal cell viability. Despite the limited SSC numbers present in BM aspirate, the current study demonstrates the advantage of SNA specificity for mRNA detection and enrichment to 1 in 200 SSCs isolated from BM aspirates. This corresponds to a 50 – 500 fold enrichment, which in together with the speed and simplicity afforded of such a strategy, demonstrates the potential of this method for broader applications.

## RESULTS AND DISCUSSION

**Scheme 1** shows a schematic illustration of the experimental approach. Cells isolated from human BMSCs directly taken from patients were incubated with SNAs for the detection of specific mRNAs. For this study two mRNA targets were chosen, runx2 and hspa8. Runx2 is a bone specific transcription factor critical in osteogenic differentiation and bone formation. Runx2 promotes the expression of osteogenesis related genes, regulates cell cycle progression and can improve the bone microenvironment.^41–43^ In contrast, hspa8 was selected as a target since the hspa8 gene encodes for the STRO-1 antigen.^44, 45^ STRO-1 is expressed on the surface of skeletal stem and progenitor populations and has been routinely used as a marker for SSC analysis and SSC sorting.^34, 40^ STRO-1 has shown to provide enrichment specificity for SSCs, as STRO-1 positive cell populations display enhanced osteogenic differentiation both *in vitro* and *in vivo*.^45^ A scrambled control sequence designed to not detect any cellular mRNA was used to demonstrate the specificity of our SNAs towards the targeted detection of mRNA. Following a 1 h incubation period of SNAs with BMSCs, FACS was used to separate target positive from target negative cells based on their fluorescence signature. Target positive cells were subsequently plated and stained to determine the number of colonies formed. SSCs have the ability to form colonies, termed colony forming units fibroblastic cells (CFU-F), when plated and cultured at limiting dilutions.^40^ The CFU-F assay harnesses the ability to identify SSCs and progenitor cells, as each colony is derived from a single stem/early progenitor cell. SSCs display upon CFU-F formation, a fibroblastic appearance when isolated and cultured.^46^

**Scheme 1.**
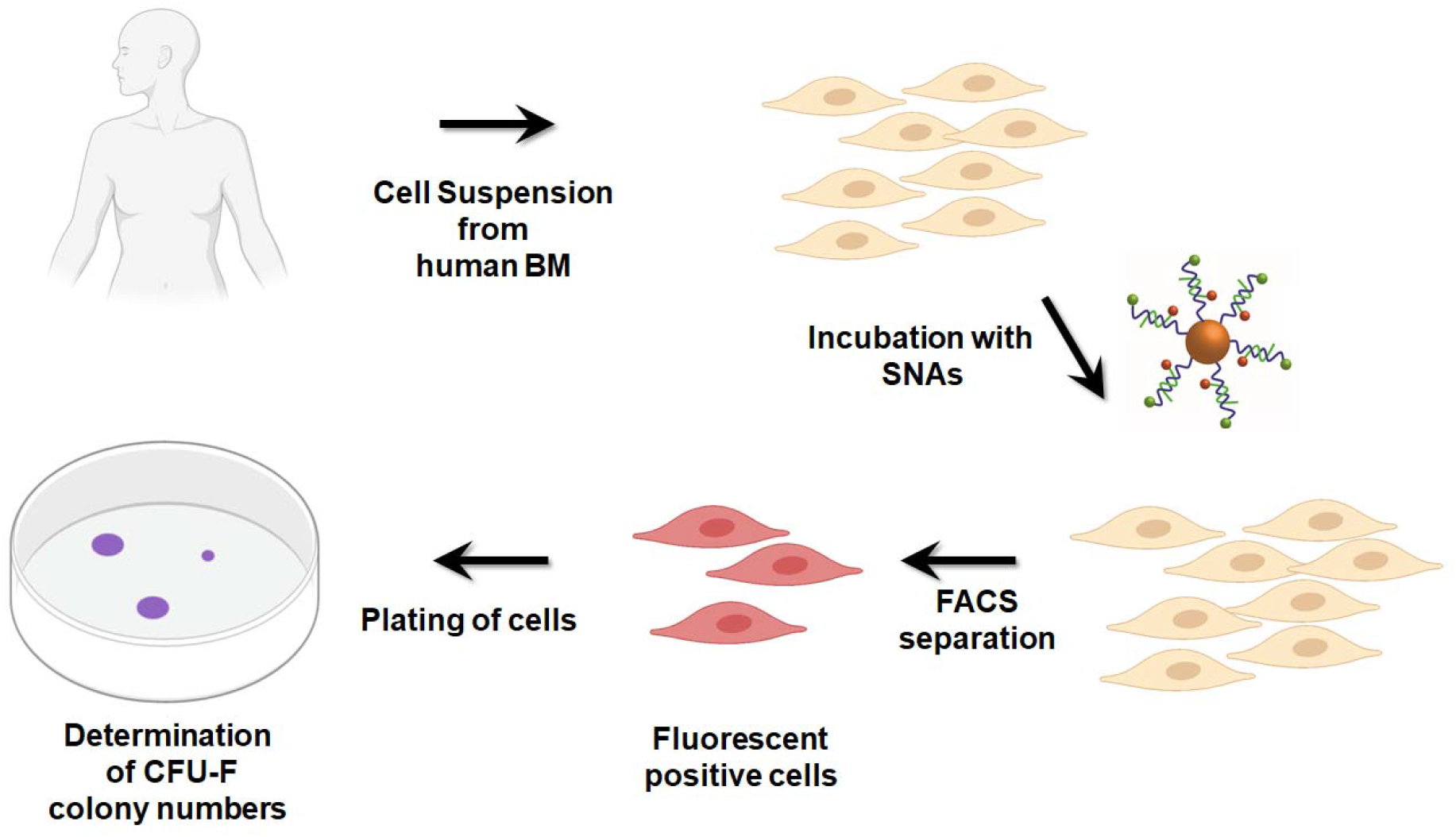
Schematic illustration of the isolation of skeletal stem cells and the formation of their colonies. Primary human bone marrow stromal cells in suspension were incubated with SNAs designed to detect hspa8 and runx2 mRNA. Following FACS separation, fluorescent cells were isolated and any SSCs formed colonies.

### Design of spherical nucleic acids for mRNA detection

**Scheme 2** demonstrates the design of SNAs for mRNA detection. Spherical AuNPs with a size of ~13 nm were functionalized with synthetic oligonucleotides bearing a terminal 3’ thiol modification and a terminal 5’ FAM dye (for simplicity the oligonucleotides directly attached to the AuNP surface are termed ‘sense’ strands). Sense strands were partially hybridized to shorter oligonucleotide complements modified with a Cy5 dye at their 5′ end (termed ‘flare’ strands). Critically, the sequence of the sense strand was designed to bind only the mRNA of interest. When both the sense and flare oligonucleotide were bound to the AuNP core, the oligonucleotide dyes were quenched (OFF state), given their close proximity to the AuNP surface. However, once the target mRNA bound to the corresponding sense sequence, the resulting displacement of the flare was detected as an increase in fluorescence at the specific wavelength of the Cy5 fluorophore (ON state). The fluorophore on the flare strand acted as a reporter, while the FAM dye on the sense strand was introduced to monitor the integrity of the sense strand, with the FAM dye released, and detectable, only if the sense strand was degraded by nucleases or detached from the nanoparticle surface.

**Scheme 2.**
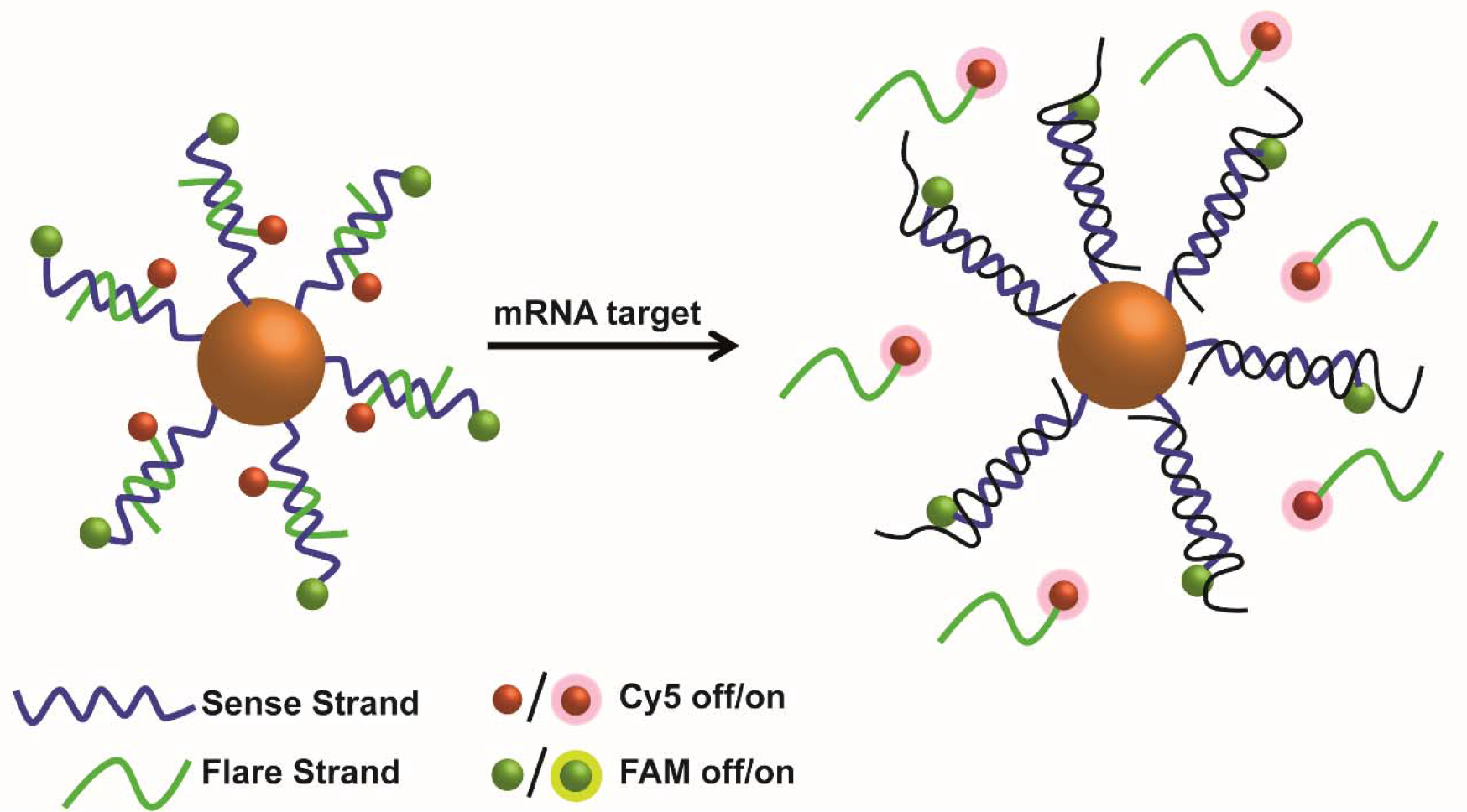
Schematic illustration of SNAs that detect mRNA. When both sense and flare strands are in close proximity to the AuNP core fluorescence is quenched. When the target mRNA is present, competitive hybridization to the sense strand leads to flare displacement and the concomitant restoration of its fluorescence signature, which is observed *via* FACS.

In this study, the synthesis of SNAs was customized for the detection of bone cell mRNAs (see **Table S1** for oligonucleotide sequences). By adopting a gradual salt aging approach ~13 nm Au NPs (see **Figure S1)** were functionalized with a shell of synthetic oligonucleotides. A shift of 4 nm was observed at the visible spectrum of the NPs indicating the change of the refractive index following attachment of the oligonucleotides (see **Figure S2**) whilst ζ – potential measurements showed that the oligonucleotide coated nanoparticles had a strong net negative charge (see **Figure S3**). Through the dissolution of the Au NP core using a solution of KI/I_2_ and the quantitative analysis of the oligonucleotides in solution, it was determined that each Au NP was coated with approximately 110 ± 4 oligonucleotide strands with no significant variation between oligonucleotides of varying sequences (see **Table S2**). Respectively, each spherical nucleic acid for the detection of mRNA consisted of approximately × 60 flare strands as shown in **Table S2**. Binding studies were also performed in a tube using perfectly matched synthetic oligonucleotide targets for each SNA (see **Figure S5**) together with stability tests, where the resistance towards nucleases was determined (see **Figure S6, S7 and S8**).

### Spherical nucleic acids for skeletal stem cell isolation and enrichment

SNAs for the detection of runx2 and hspa8 mRNA were incubated with human BMSCs for 1, 2 or 3 h. Following determination of the appropriate settings for FACS separation (see **Figure S9**), the fluorescence signal corresponding to mRNA detection and the subsequent flare release was monitored by observing an increase in Cy5 fluorescence in comparison to cells incubated with no SNAs. **Figure 1** shows that following addition of runx2 and hspa8 SNAs, Cy5 positive cells could be detected within 1 h of incubation in the sub-population of BM cells including lymphocytes, granulocytes and monocytes, while the fluorescence intensity was observed to increase with longer incubation times (2 – 3 h). Moreover, the absence of a signal from the FAM dye conjugated to the sense strand indicated the absence of sense strand degradation (**Figure S10**). In both cases, a scrambled SNA designed to not detect any intracellular mRNA was also employed as a control. The absence of both a Cy5 and FAM signal from the flare and sense strand respectively, after a 3 h incubation period, indicated not only strand stability against nuclease degradation but also the specificity of SNAs for the detection of mRNA.

**Figure 1.**
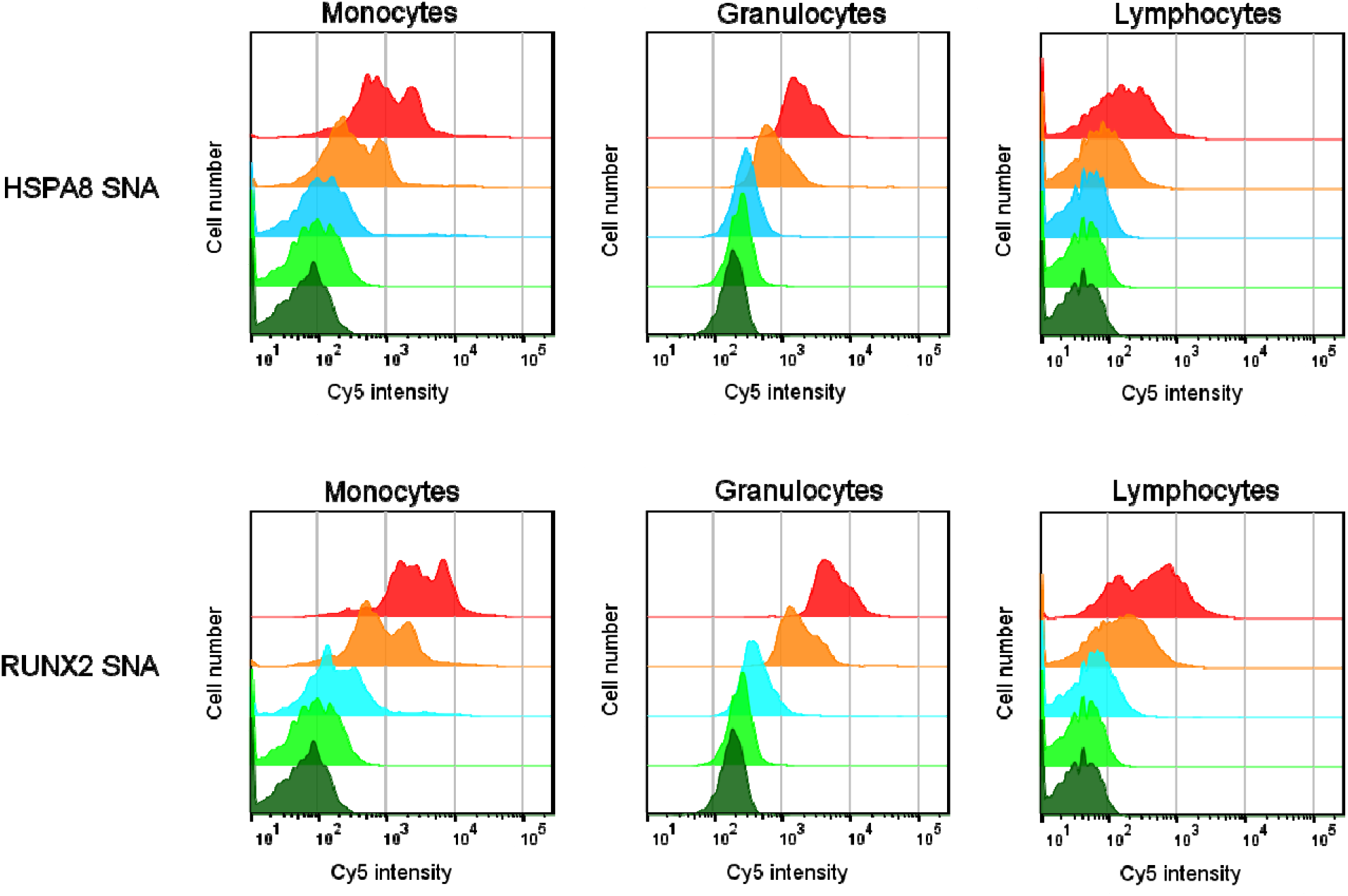
SNAs for the detection of hspa8 and runx2 mRNA including a scrambled SNA were incubated with human bone marrow stromal cells for 1, 2 or 3 hours at 0.2 nM final concentration. After washing, the cells were analyzed by flow cytometry for Cy5 fluorescence intensity. Cy5 intensity versus cell number overlay plots for the different SNAs for different cell types within the bone marrow as determined by the FACS selection criteria. Color guide: Red – 3 hour incubation, Orange – 2 hour incubation, Light blue – 1 hour incubation, Light green – scrambled control at 3 hour incubation, Dark green – no SNAs at 3 hour incubation.

Following cell sorting of Cy5 positive cells for runx2 and hspa8 mRNA, the recovered cells were grown for 14 days then stained, and the number of colonies formed determined. Only cells of monocyte size and granularity were collected by FACS, as this corresponds to SSC characteristics, whilst other cell types were discarded.^47, 48^

**Figure 2A** shows that isolated cells produced varying levels of CFU-Fs, with cells sorted from SNAs designed for the detection of runx2 mRNA producing a higher number of cell colonies than hspa8 with significant enrichment observed in comparison to unsorted cells (see **Figure S11**). In contrast, the fraction of non-fluorescent cells (Cy5 negative) showed no/negligible CFU-F formation (see **Table S3** for patient specific CFU-F values). **Figure 2B and C** shows that cells isolated using the SNAs for the detection of hspa8 (**B**) and runx2 (**C**) mRNA displayed a fibroblastic phenotype. Through the collection of different proportions of the brightest Cy5 cells (15 – 60 %) with the two different SNAs, regardless of their overall intensity, it was demonstrated CFU-Fs occurred in the top 15 % of brightest cells for these two targets.

**Figure 2.**
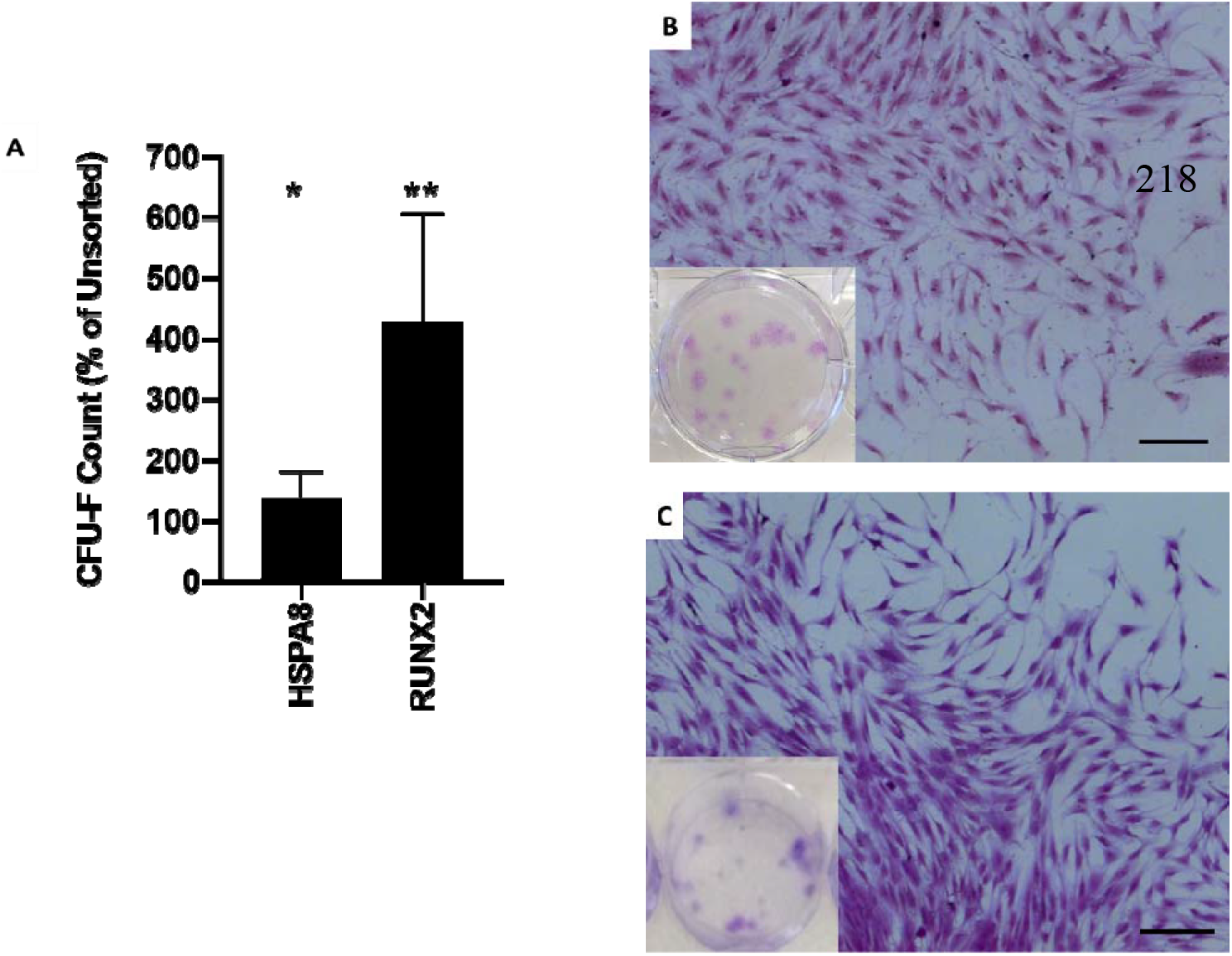
(A) Graph shows CFU – F count for sorted cells as a percentage of the CFU – F number from unselected cells which were plated to give the equivalent of 10,000 cells in the monocyte region per well. Graphs show means with SD. n=3 patients for each SNA. * p<0.05, ** p>0.01. (B, C) Representative images of cells within individual CFU-F colonies isolated with SNAs for the detection of hspa8 (A) and runx2 (B) mRNA. Cells isolated using SNAs displayed a fibroblastic phenotype, characteristic of such colonies. Insets show individual colonies in a 6-well plate. Scale bar is 200 μm.

To determine if the cells isolated were indeed SSCs with tri-lineage potential and the capacity to generate the requisite stromal lineages, the collected cells, isolated using the runx2 SNA, were grown and expanded and then subjected to different culture conditions to generate osteogenic, adipogenic and chondrogenic stromal populations. **Figure 3** demonstrates that cells maintained under osteogenic conditions produced enhanced positive staining for alkaline phosphatase (see **Figure 3A**) compared to basal conditions. Alcian blue staining of the chondrogenic pellets revealed proteoglycan synthesis (see **Figure 3B**) whereas Sirius Red stain retention was absent in pellets, indicating an absence of collagenous matrix formation. Oil-Red-O staining of lipids provided evidence of adipogenesis and lipid droplet formation (see **Figure 3C**), with no lipid visible in basal conditions. The results demonstrate that the runx2 SNA-isolated cells displayed tri-lineage potential and generated stromal populations with osteogenic, adipogenic and chondrogenic potential thus indicating that the isolated cells contained SSCs.

**Figure 3.**
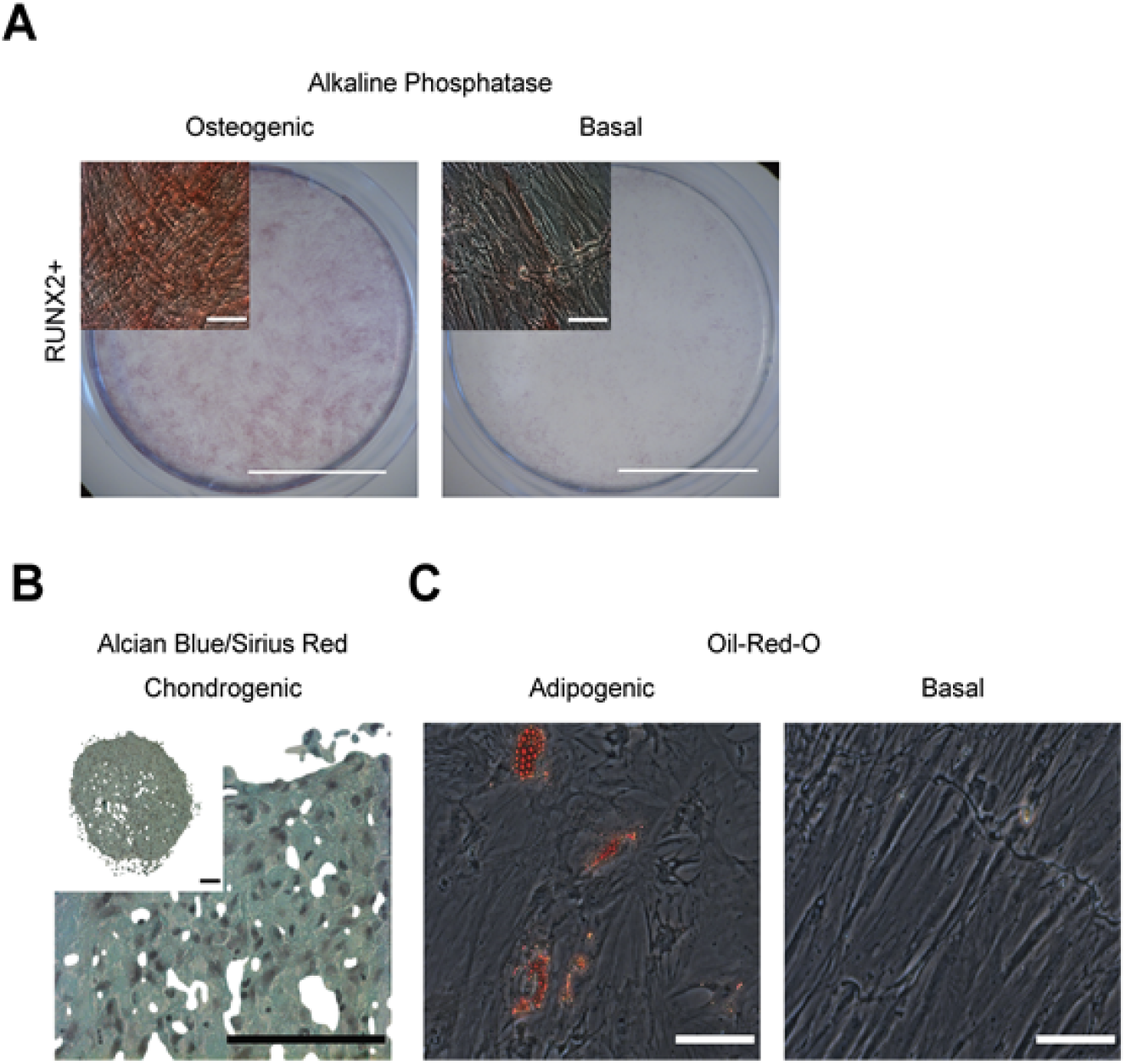
Cells isolated using the runx2 SNA display capacity for tri-lineage stromal cell differentiation. Isolated populations demonstrated (A) osteogenic, (B) chondrogenic and (C) adipogenic induction indicating the presence of SSCs.

## CONCLUSION

The current study demonstrates the development of SNAs for the detection of specific mRNA targets, runx2 and hspa8, in human BM stromal populations, which enables the isolation and enrichment of human SSCs. Following a short incubation period with runx2 and hspa8 SNAs with human BMSCs, fluorescent cells were sorted using FACS. Further plating of isolated mRNA target positive cells resulted in a significant enrichment in CFU-F formation. Using this technique, we demonstrate that despite the low abundance of SSCs in human BM, up to 1 in 200 of the Cy5 positive cells isolated by FACS formed colonies with high CFU-F levels on culture.

In conclusion, the current study demonstrates the potential of SNAs for the isolation and enrichment of skeletal stem cell populations with exciting opportunities therein for translational application, one of our future research goals.

## EXPERIMENTAL METHODS

### SNAs for mRNA Detection

Briefly bis(p-sulfonatophenyl)phenylphosphine dihydrate dipotassium salt (BSPP) coated 13 nm AuNPs (10 nM, 1 mL) synthesised using the Turkevich method were incubated with thiol – modified synthetic oligonucleotides (3 μM, 1 mL) (see **Table S1** for oligonucleotide sequences).^49^ After the mixture was left shaking for 24 h, BSPP (1mg/ 20 μL, 10 μL), phosphate buffer (0.1 M, pH 7.4) and sodium dodecyl sulphate (SDS, 10 % solution) were added to achieve a final concentration of 0.01 M phosphate and 1 % SDS respectively. A gradual salt aging was then performed by six equal additions of NaCl (2 M) over 8 h in order to achieve a final salt concentration of 0.3 M in solution. After shaking overnight, the resulting SNAs were purified by three rounds of centrifugation (16,400 rpm, 20 min) and were stored at 4 °C in phosphate buffer saline (PBS).

For flare hybridization, SNAs (16 nM, 500 μL) were mixed with an excess of complementary flare oligonucleotides (960 nM, 500 μL). The solution was heated to 60 °C for 5 min, followed by slow cooling to room temperature. Samples were purified by two rounds of centrifugation (16,400 rpm, 15 min) and re – dispersed in PBS.

### Incubation of SNAs with BMSCs in suspension

The SNAs were added to the BMSC suspension (at 10^6^ cells per mL) to the required final concentration of 0.1 or 0.2 nM and for the appropriate incubation time. Cells were subsequently washed in basal medium and then resuspended in FACS solution (0.5% BSA, 2mM EDTA in 1x PBS) prior to FACS analysis. The FACS Aria cytometer (Becton Dickinson, Wokingham, UK) was used to acquire 20000 cells with data analysed using the FlowJo software version 10.6.1. All washing stages were at 400 x g for 5 minutes.

### Processing of human BM samples

Bone marrow samples were obtained from haematologically healthy patients undergoing hip replacement surgery with local ethics committee approval (LREC194/99/1 and 18/NW/0231 and 210/01) and informed patient consent. In brief, bone marrow was first washed at least 3 times in 50 mL α – MEM medium to remove fat, then passed through a 70 μm cell strainer. Marrow cells were resuspended in 10 mL α – MEM and subjected to density centrifugation using 20 mL Lymphoprep™ (Lonza) at 800 x g for 20 minutes with no braking on the centrifuge. The buffy coat layer, containing bone marrow mononuclear cells was washed in 5 ml basal medium (α **–** MEM containing 10% FBS and 100 U mL^−1^ penicillin and 100 μg mL^−1^ streptomycin; Lonza). All washing stages were at 400 x g for 5 minutes.

### Colony Forming Units – Fibroblast (CFU-F) Assay

A FACS Aria was used to collect SNA positive and negative cells from human bone marrow samples. Cells were gated for monocytes, single cells, and Cy5 fluorescence. Positive samples were deemed to be the top 15% of Cy5 fluorescent cells and negative samples were deemed to be the lower 15% of Cy5 cells collected. Cy5 positive and negative cells were collected by FACS sorting. Ten thousand cells were placed into each well of 6 – well tissue culture plates containing 2 ml basal medium. Cells were grown for 14 days, with a medium change after 7 days. On day 14, wells were washed with 3 ml PBS and then fixed with 1 mL 95% ethanol for 10 minutes. The wells were subsequently air dried and 1 mL 0.05% crystal violet solution added to each well for 1 minute. The wells were then washed twice with 2 mL distilled water and the number of visible colonies determined by eye.

### Osteogenic Differentiation Assay

Passage 1 cells were cultured at 37 °C in 5% CO_2_ until confluent, then seeded at 10,000 cells per well on a 12 well plate followed by culture in basal media for 24 hours. Cells were then cultured in osteoinductive media (basal medium with 50 μM ascorbic acid 2-phosphate and 10 nM vitamin D_3_) for 14 days at 37 °C in 5% CO_2_ with media change every 3-4 days. Cells were washed in PBS, fixed in 95% ETOH, then stained with alkaline phosphatase.

### Adipogenic Differentiation Assay

Passage 1 cells were cultured at 37 °C in 5% CO_2_ until confluent, then seeded at 10,000 cells per well on a 12 well plate followed by culture in basal media until 80% confluent, after approximately 3-5 days. Cells were then cultured in adipogenic media (basal medium with 100 nM dexamethasone, 500 μM IBMX, 3 μg/ml ITS solution, and 1 μM rosiglitazone) for 14 days at 37 °C in 5% CO_2_ with media change every 3-4 days. Cells were washed in PBS, fixed in 4% paraformaldehyde, washed in PBS again and then stained with Oil Red O.

### Chondrogenic Differentiation Assay

Passage 2 cells were cultured at 37 °C in 5% CO_2_ until confluent, then diluted to 500,000 cells per ml in chondrogenic media (α-MEM containing 100 U mL^−1^ penicillin and 100 μg mL^−1^ streptomycin, 100 μM ascorbic acid 2-phosphate, 10 ng/ml TGF-B_3_, 10 μg/ml ITS solution, 10 nM dexamethasone) in a universal container. Cells were centrifuged at 400 x g for 10 minutes to form a cell pellet, and all but 1 ml of media removed. Cells were then cultured with the tube cap loose for 14 days at 37 °C in 5% CO_2_ with media change every 2 days. Cells were washed in PBS, fixed in 95% ETOH, then stained with alcian blue and Sirius red.

### Statistical Analysis

Wilcoxon – Mann – Whitney statistical analysis and ANOVA were performed where appropriate using the SPSS for Windows program version 23 (IBM Corp, Portsmouth, Hampshire, UK). All experiments were completed at least three times and data are presented as mean ± SD. Significance was determined with a p – level of 0.05 or lower.

## ASSOCIATED CONTENT

The Supporting Information is available free of charge on the ACS Publications website at DOI: XXXX Additional experimental detail and data including SNAs additional characterization (UV-Vis, zeta potential, TEM), oligonucleotide sequences, FACS settings, and CFU-F counts. The raw data is available at https://doi.org/10.5258/SOTON/D1440

## Notes

The authors declare no competing financial interest.

## ACKNOWLEDGEMENTS

ROCO and AGK acknowledge financial support from the Biotechnology and Biological Sciences Research Council (BB/P017711/1). ROCO acknowledges support from the UK Regenerative Medicine Platform “Acellular / Smart Materials – 3D Architecture” (MR/R015651/1), the Rosetrees Trust and Wessex Medical Research.

