## Supplementary data for "Enrichment of Skeletal Stem Cells from Human Bone Marrow Using Spherical Nucleic Acids"

**Table of Contents**

**S – I	SNA Characterization**

a. **Transmission Electron Microscopy (TEM)**
b. **UV** – **Vis spectroscopy**
c. **ζ – potential**
d. **Design of oligonucleotide sequences**
e. **Synthesis of oligonucleotides**
f. **Quantitative assessment of oligonucleotide loading**
g. **Background fluorescence spectra**
h. **SNA response to complementary ssDNA**
i. **SNA nuclease stability**

**S – II	Supplementary Experimental Data**

a. **Determination of settings for FACS separation**
b. **Stability of SNAs against nuclease degradation**
c. **CFU** – **F counts for SNAs**

**S – III References**

**S – I	SNA Characterization**

a. **Transmission Electron Microscopy (TEM)**

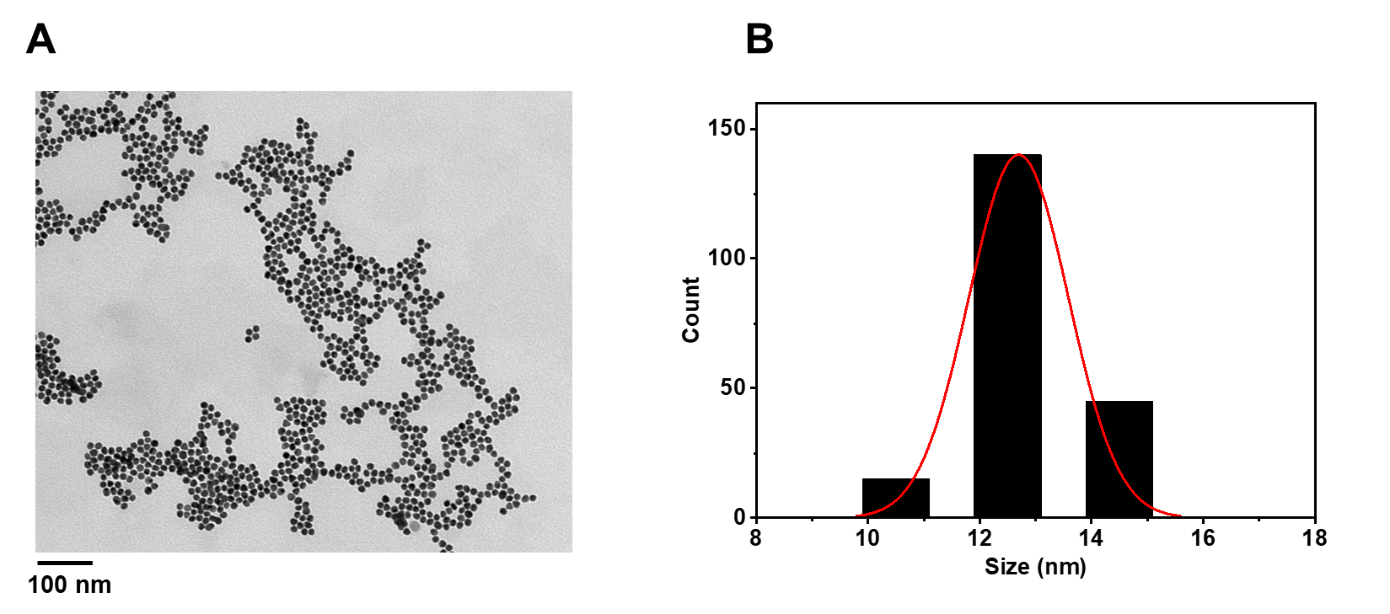

**Figure S1.** A) TEM image of bis(p-sulfonatophenyl)phenylphosphine dihydrate dipotassium salt (BSPP) – coated AuNPs dispersed in Milli – Q water and deposited on a copper grid. B) Corresponding size distribution histogram. Average AuNP size was determined by calculating the size of 200 AuNPs *via* imageJ.

b. **UV** – **Vis spectroscopy**

Following functionalization and purification, the visible spectra of SNAs were acquired and compared to BSPP functionalized AuNPs. A clear shift was observed following DNA attachment, which is due to a change in the refractive index.

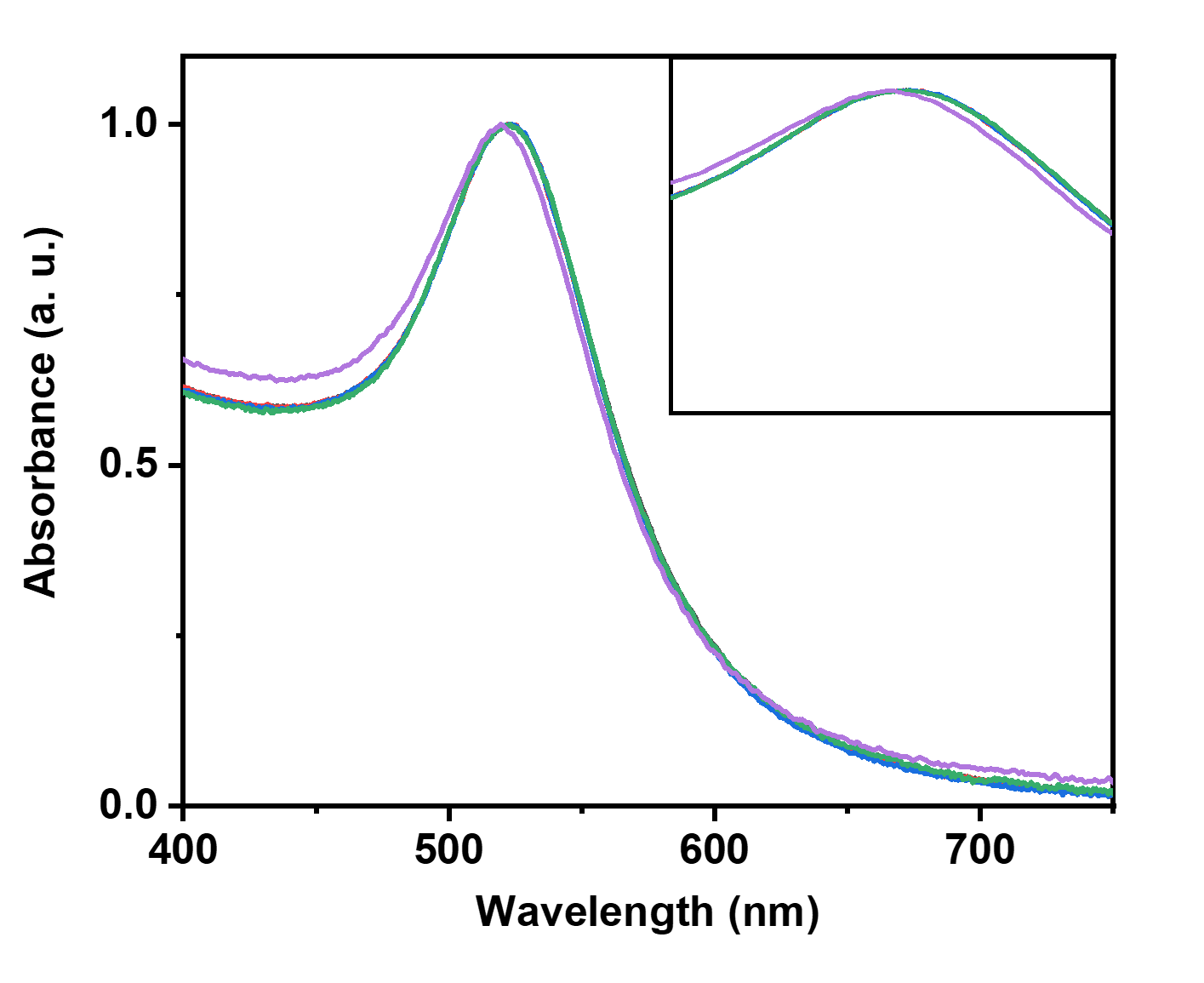

**Figure S2.** Normalized visible spectra of SNAs. A 4 nm shift was observed after oligonucleotide functionalization indicating a change in the refractive index and successful surface functionalization. Colour Guide: Black – hspa8, Red – runx2, Blue – scramble, Green – vimentin, Purple – BSPP coated AuNPs

c. **ζ – potential**

The net surface charge of BSPP and SNAs was assessed *via* ζ – potential.

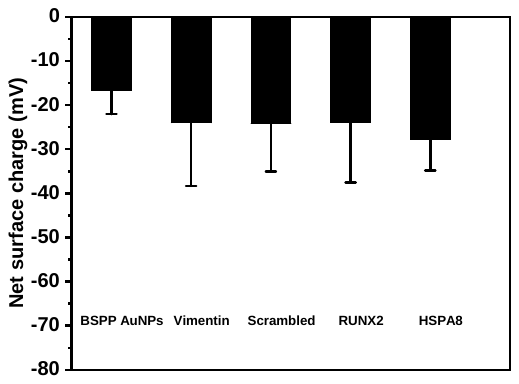

**Figure S3.** Net surface charge of BSPP AuNPs and SNAs. After ligand replacement of BSPP with thiol modified oligonucleotides a decrease in the net surface charge was observed as a result of the negatively charged oligonucleotides.

d. **Design of oligonucleotide sequences**

Appropriate sequences were determined using the Basic Local Alignment Sense Tool (BLAST) using the following settings: nucleotide-specific blast (blastn), database: RefSeq mRNA, Species: Homo Sapien, Expected Threshold: 10 Match/Mismatch scores: 2/-3. A sense sequence was designed to have a length of 21-23 bases, a GC content ˂ 50 %, an E value ˂ 0.05 and an E value of nearest match > 1. A flare strand was designed to show partial complementarity to the sense sequence with a length of 10-12 bases and a melting temperature of > 40 ºC.

**Table S1.** Sense and flare oligonucleotide sequences. X: thiol modifier C3 S – S (CPG resin from Glen Research)

| **DNA – coated AuNPs** | **Oligonucleotide sequences (5’ to 3’) and modifications** |
| --- | --- |
| hspa8 (sense strand)  hspa8 (flare strand) | FAM – AGCAGTACGGAGGCGTCTTACAAAAAAAAA – X  Cy5 – TGTAAGACGCCTC |
| runx2 (sense strand)  runx2 (flare strand) | FAM – TGTGGTTGTTTGTGAGGCGAATAAAAAAAA – X  Cy5 – ATTCGCCTCACA |
| scrambled control (sense strand)  scramble control (flare strand) | FAM – ATGGTATACCGAAAGACTGTTAAAAA – X  Cy5 – AACAGTCTTTCG |
| vimentin (sense strand)  vimentin (flare strand) | FAM – CTTTGCTCGAATGTGCGGACTTAAAAAAAA – X  Cy5 – AAGTCCGCACA |

e. **Synthesis of oligonucleotides**

Oligonucleotides were synthesized on an Applied Biosystems 394 automated DNA/RNA synthesizer using a standard 1.0 μmole phosphoramidite cycle of acid-catalyzed detritylation, coupling, capping, and iodine oxidation. Stepwise coupling efficiencies and overall yields were determined by the automated trityl cation conductivity monitoring facility and was >98.0%. Solid support 3'-Thiol-Modifier C3 S-S CPG, 1000/110, item number: 2361, 5'-Fluorescein-CE Phosphoramidite (6-FAM), item number: 2134 and Cy-5-CE phosphoramidite (Cyanine 650) item number 2521 were purchased from Link Technologies Ltd. Additional reagents were purchased from Applied Biosystems Ltd. All β-cyanoethyl phosphoramidite monomers were dissolved in anhydrous acetonitrile to a concentration of 0.1 M immediately prior to use with coupling time of 50 s for normal A, G, C, and T monomers and was extended to 600 s for modified monomers. Cleavage and deprotection were achieved by exposure to concentrated aqueous ammonia solution for 60 min at room temperature followed by heating in a sealed tube for 5 h at 55 °C. Purification was carried out by reversed-phase HPLC on a Gilson system using a Brownlee Aquapore column (C8, 8 mm x 250 mm, 300Å pore) with a gradient of acetonitrile in triethylammonium bicarbonate (TEAB) increasing from 0% to 50% buffer B over 30 min with a flow rate of 4 mL/min (buffer A: 0.1 M triethylammonium bicarbonate, pH 7.0, buffer B: 0.1 M triethylammonium bicarbonate, pH 7.0 with 50% acetonitrile). Elution was monitored by ultraviolet absorption at 298 nm. After HPLC purification, oligonucleotides were freeze dried then dissolved in water without the need for desalting. All Purified oligonucleotides were characterised by electrospray mass spectrometry. Mass spectra of oligonucleotides were recorded either using a Bruker micrOTOFTM II focus ESI-TOF MS instrument in ES- mode or a XEVO G2-QTOF MS instrument in ES- mode. Data were processed using MaxEnt and in all cases confirmed the integrity of the sequences.

f. **Quantitative assessment of oligonucleotide loading**

**Table S2.** Determination of ‘sense’ and ‘flare’ strand oligonucleotide loading for each type of SNA. Values based on triplicate experiments.

| **Strand** | **Average number of oligonucleotides / AuNP** | **Standard error** |
| --- | --- | --- |
| runx2 sense strand  runx2 flare strand | 109  59 | 4  2 |
| hspa8 sense strand  hspa8 flare strand | 111  58 | 4  1 |
| scramble sense strand  scramble flare strand | 109  61 | 5  2 |
| vimentin sense strand  vimentin flare strand | 114  61 | 6  2 |

g. **Background fluorescence spectra**

The fluorescence intensity of the dyes associated with the SNAs was evaluated in order to assess the degree of background fluorescence resulting from incomplete quenching of the Cy5 and FAM dyes.

Fully assembled SNAs (6.7 nM, 150 μL) in PBS were left at 37 °C for 24 h after which the fluorescence intensities of Cy5 (flare strand dye) and FAM (sense strand dye) were evaluated in a tube. **Figure S4** shows that in each case the background fluorescence of both dyes is minimal with no significant variation in intensity between SNAs.

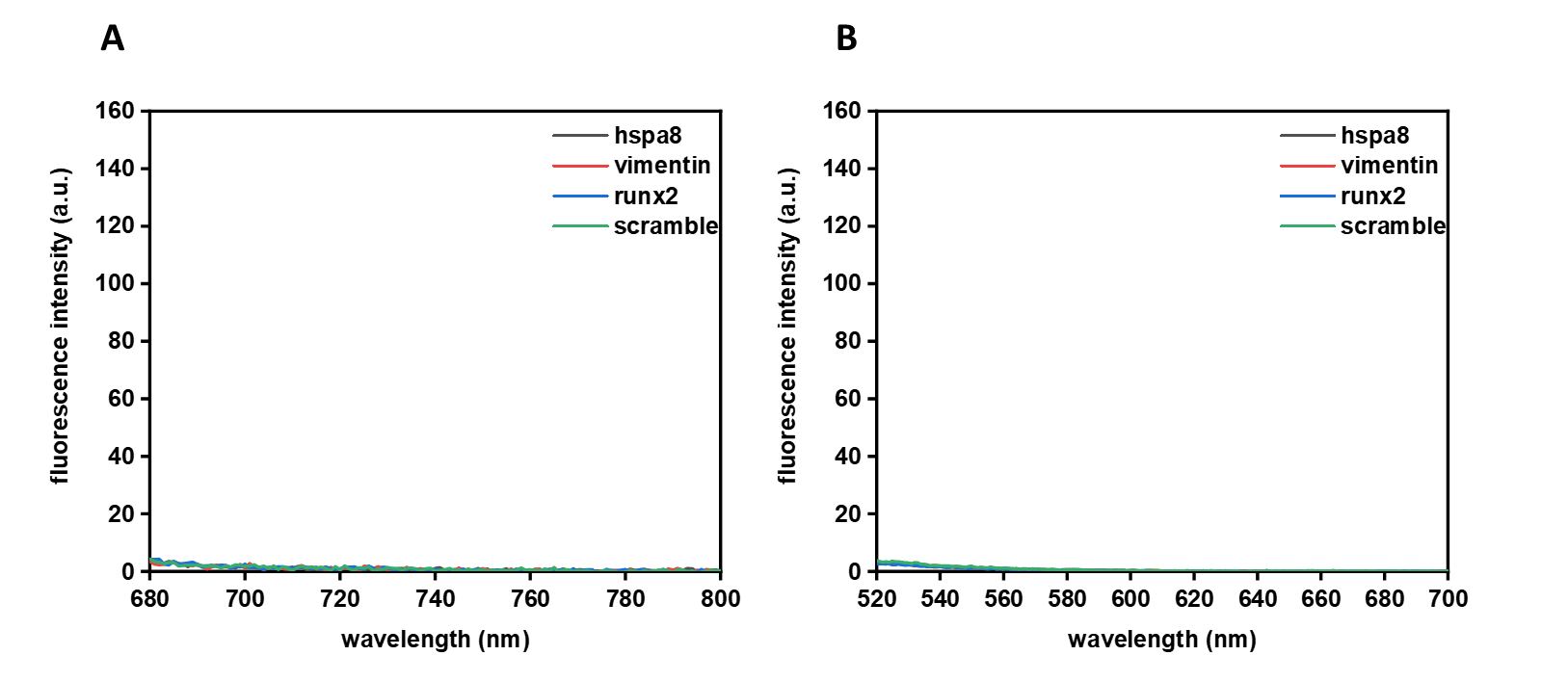

**Figure S4**. Background fluorescence spectra of Cy5 (A) and FAM (B) dyes for each SNA. Spectra were acquired on a Carry Eclipse Fluorescence Spectrophotometer.

h. **SNA response to complementary ssDNA**

SNAs for mRNA detection were tested with synthetic fully complementary oligonucleotide targets. SNAs (6.7 nM, 150 μL) in PBS were incubated with their fully complementary target. **Figure S5** shows that in the presence of the fully complementary target SNAs responded with an increase in the Cy5 fluorescence signal upon recognition and binding within the first 2 minutes indicating their efficiency at signaling the presence of a specific target in buffered conditions.

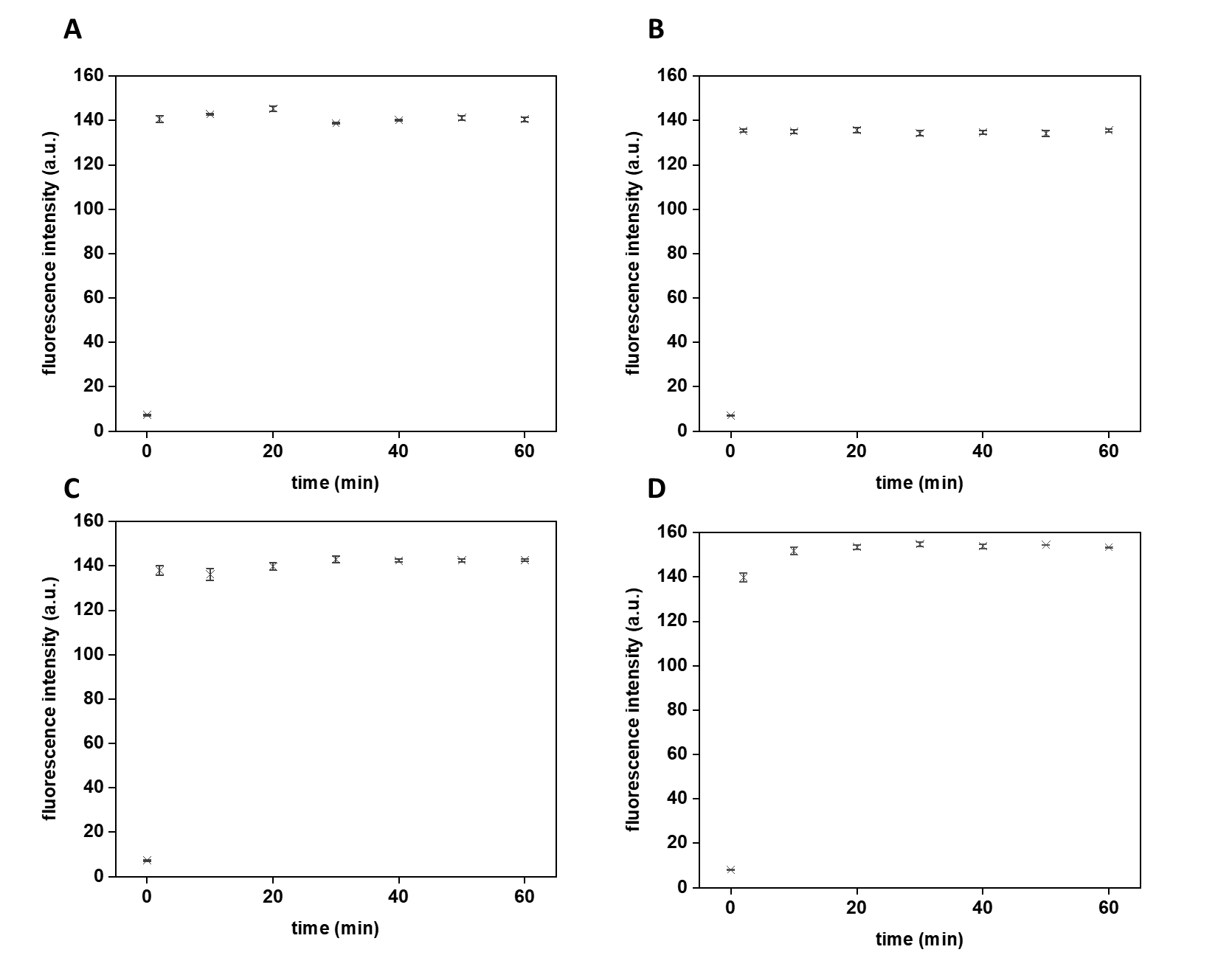

**Figure S5**. Time course of fluorescence associated with flare release when SNAs were incubated with synthetic oligonucleotide strands leading to the displacement of the hspa8 (A), vimentin (B), runx2 (C) and scramble (D) flare.

i. **SNA nuclease stability**

Stability of SNAs was investigated in the presence of DNase I. Prior to the experiment the activity of the enzyme was assessed. Initially, a free duplex of DNA was incubated with DNaseI and visualized using polyacrylamide gel electrophoresis (PAGE) in order to assess the efficiency of DNaseI to degrade free dsDNA. **Figure S6** shows that after a 24 h incubation at 37 °C the duplex was fully degraded indicating efficient enzyme activity.

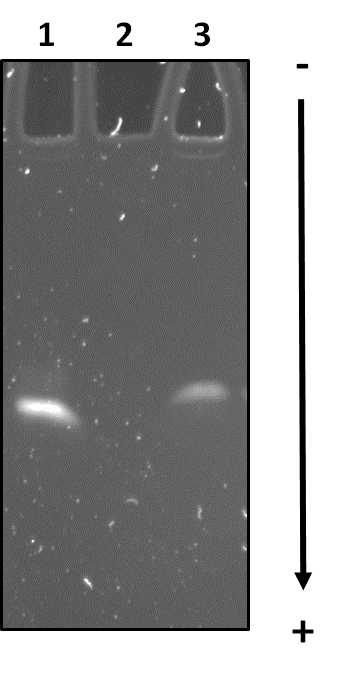

**Figure S6.** PAGE gel displaying ssDNA (lane 1), dsDNA incubated with DNaseI for 24 h (lane 2) and dsDNA not incubated with DNaseI (lane 3). The disappearance of a clear band in lane 2 indicated that DNaseI was active in the presence of a free duplex.

SNAs (2.5 nM, 150 μL) in a DNase I buffer containing Tris-HCl (10 mM), MgCl_2_ (2.5 mM) and CaCl_2_ (0.5 mM) at pH 7.4 were then incubated with DNase I (from bovine pancreas, Sigma Aldrich, 2 U/L). Samples were then incubated at 37 °C for 24 h. By monitoring the fluorescence intensity of the sense strand (FAM), the % of oligonucleotides remaining on the AuNP was determined. **Figure S7** shows that in the presence of DNase I, SNAs retained more than 95 % of oligonucleotide surface coating suggesting that our SNAs displayed unique stability.

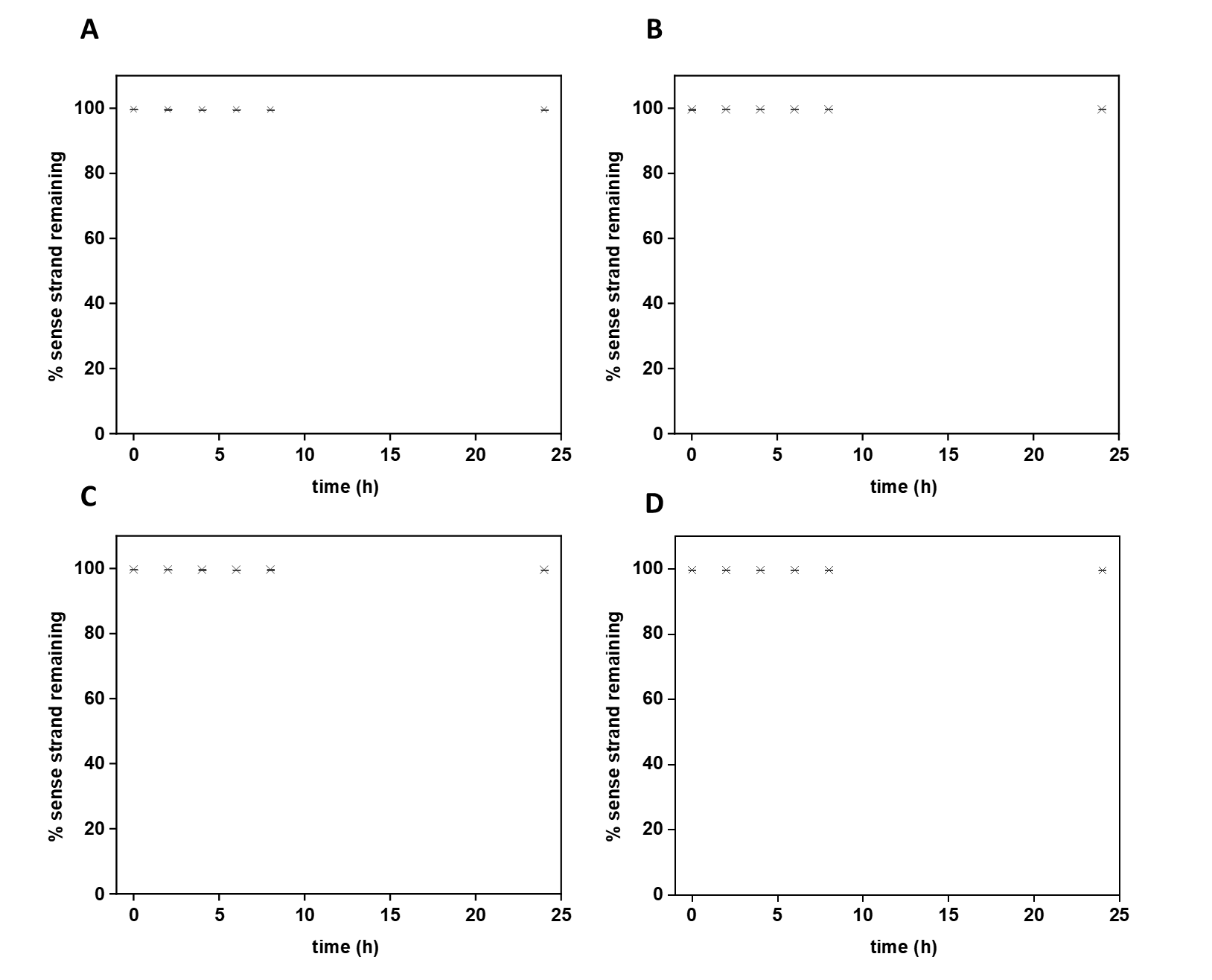

**Figure S7**. SNAs for the detection of hspa8 (A), vimentin (B) and runx2 (C) mRNA including a scramble SNA (D) were incubated with DNase I over a 24 h period. By monitoring the fluorescence intensity, the % of oligonucleotides on the AuNP surface was determined.

A suggested factor for the increased resistance within cells is the high local salt concentration at the NP surface.^1-4^ Previous work has demonstrated that monovalent cations, including Na^+^, inhibit DNase I as well as related nucleases. This inhibition is caused by the displacement of Ca^2+^ and Mg^2+^ by Na^+^ ions that are bound to the enzyme and are required for efficient activity. An increased concentration of Na^+^ ions has further been associated with an increase in oligonucleotide surface density that requires more charge balancing counterions.^4^ Pellegrino *et al.* have also shown that DNA attached to a AuNP surface can undergo conformation changes.^5^ These changes can therefore prevent binding of DNase I, which is known to be a minor groove binder.^6^ Nevertheless, it can be concluded that the demonstrated resistance is in part due to the dense packing of the oligonucleotides and the resulting high cation concentration that prevents efficient enzyme activity.

We therefore attributed the increased resistance observed in our study to the dense oligonucleotide loading.^7^ In order to experimentally demonstrate the effect of oligonucleotide density on the NPs, a SNA with a lower oligonucleotide density (40 oligonucleotides/ AuNP) was subjected to the same nuclease conditions. The FAM signal from the sense strand was monitored over 24 h and compared to the signal obtained with a high density SNA used in or study **Figure S8** shows that after 2 h a clear increase in the FAM signal was observed which did not significantly change over the 24 h incubation period.

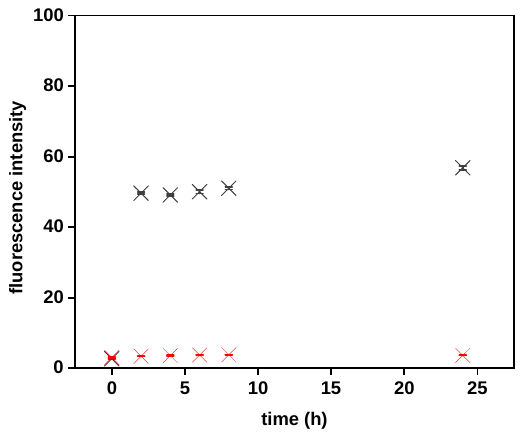

**Figure S8.** Fluorescence intensity of FAM signal monitored over 24 h of low oligonucleotide density (black) and high oligonucleotide density (red) SNAs incubated with DNaseI. A clear increase in the FAM intensity was observed when SNAs were coated with approximately 40 oligonucleotides/AuNP (low density) indicating that oligonucleotide degradation occurred. In comparison when coated with a larger number of oligonucleotides (~100) no significant degradation was observed thus proving that dense packing of oligonucleotides on the AuNP surface can prevent enzyme activity.

**S – II Supplementary Experimental Data**

a. **Determination of settings for FACS separation**

The appropriate settings for FACS separation were established. **Figure S9** shows how the flow cytometry data were gated against time to remove events occurring before the flow stream and laser power had stabilized. Debris were eliminated by gating out events with low forward (FSC **–** H) and side scatter (SSC **–** H) and doublets were excluded following a pulse geometry gating strategy by omitting cells that did not follow a linear increase of the pulse area vs height on an FSC scatter plot. Finally, sub **–** populations were gated for cells falling inside typical FSC and SSC distributions for white blood cells found in human BM including lymphocytes, granulocytes and monocytes, as shown in previous publications.^8, 9^ Lymphocytes are typically smaller cells with low FSC and granulocytes have higher SSC intensity because of their granular cytoplasmic content.^10^

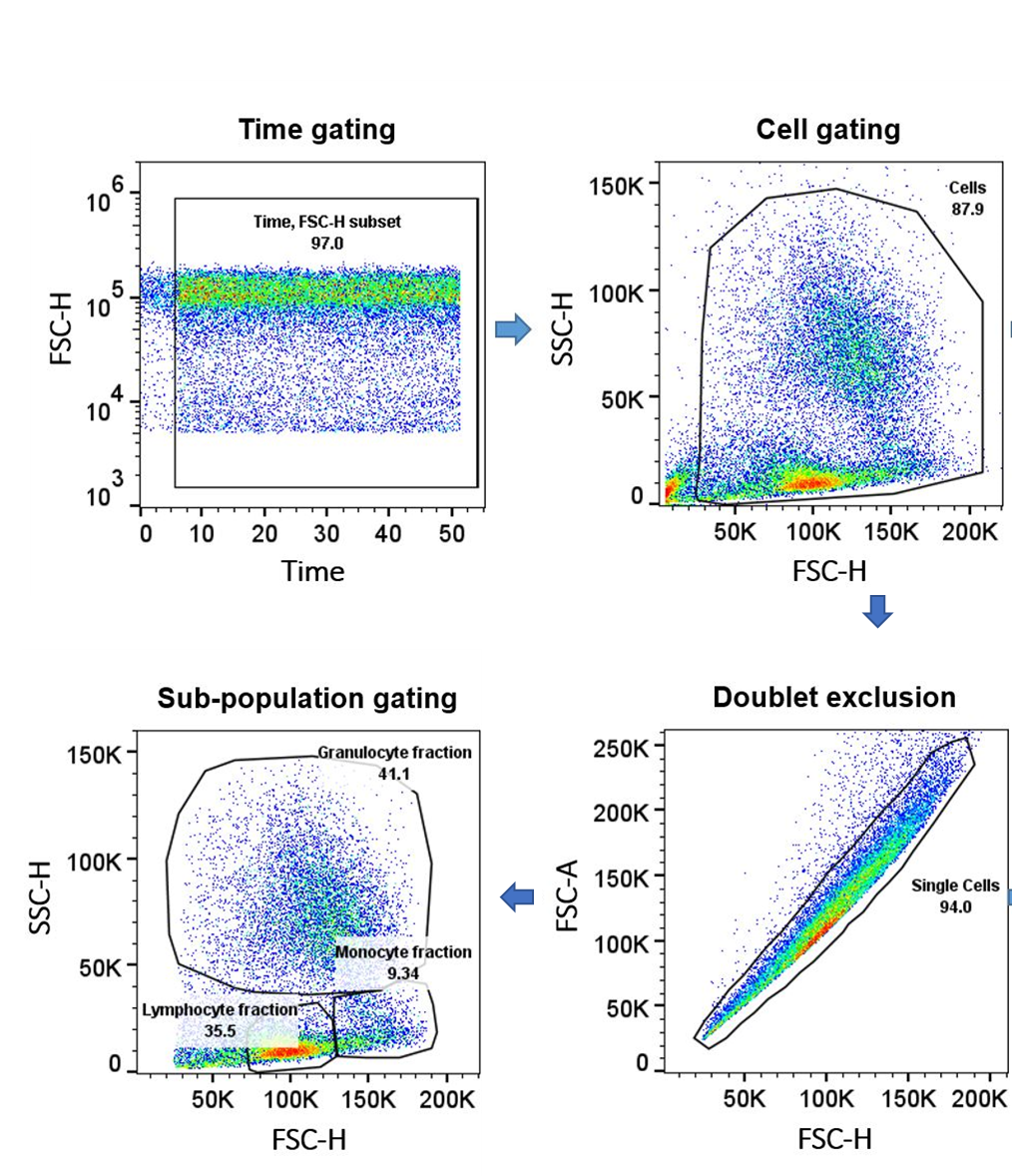

**Figure S9.** Flow cytometry selection criteria for the different cell fraction regions found in human bone marrow samples. Data were gated against time; debris were eliminated by gating out events and finally sub population of white blood cells were separated.

b. **Stability of SNAs against nuclease degradation**

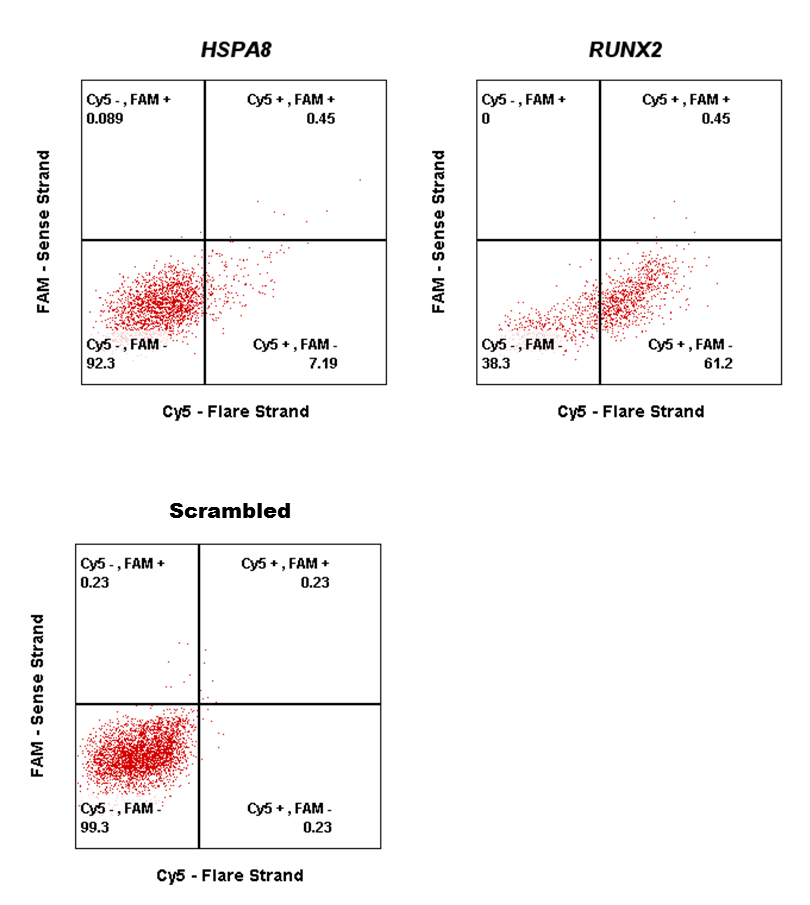

**Figure S10.** Representative FACS Cy5 vs FAM dot plots shown for human bone marrow samples used to isolate and enrich for human skeletal stem cells. The absence of any significant FAM fluorescence regardless of the target detected demonstrates that the presence of a Cy5 fluorescence is due to target detection rather than degradation. Furthermore, the absence of a fluorescence signal from both Cy5 and FAM following incubation with a scramble SNA indicates the specificity of SNAs towards the detection of mRNA.

c. **CFU-F counts for SNAs**

**Table S3.** CFU-F counts as percentage of unsorted cell CFU-F count, from Cy5 positive and Cy5 negative cells isolated from different patients using the stated SNAs.

|  | | CFU-F Count as percentage of unsorted cell CFU-F Count | | | | | | | | |
| --- | --- | --- | --- | --- | --- | --- | --- | --- | --- | --- |
| SNA  target | Cell fraction | Patient 1 | | | Patient 2 | | | Patient 3 | | |
| hspa8 | Cy5 + | 111 | 77 | 77 | 157 | 198 | 123 | 300 | 100 | 100 |
|  | Cy5 - | 0 | 0 | 0 | 0 | 0 | 0 | 0 | 0 | 0 |
| runx2 | Cy5 + | 386 | 643 | 300 | 275 | 225 | 225 | 500 | 600 | 700 |
|  | Cy5 - | 0 | 86 | 0 | 0 | 0 | 0 | 0 | 0 | 0 |

**S – III References**

9.	Xavier, M.; de Andres, M. C.; Spencer, D.; Oreffo, R. O. C.; Morgan, H., Size and dielectric properties of skeletal stem cells change critically after enrichment and expansion from human bone marrow: consequences for microfluidic cell sorting. *Journal of the Royal Society Interface* **2017,** *14* (133).

10.	Fujimoto, H.; Sakata, T.; Hamaguchi, Y.; Shiga, S.; Tohyama, K.; Ichiyama, S.; Wang, F.; Houwen, B., Flow cytometric method for enumeration and classification of reactive immature granulocyte populations. *Cytometry* **2000,** *42* (6), 371-378.
